# Dendritic cell-mediated responses to secreted *Cryptosporidium* effectors are required for parasite-specific CD8^+^ T cell responses

**DOI:** 10.1101/2023.08.16.553566

**Authors:** Breanne E. Haskins, Jodi A. Gullicksrud, Bethan A. Wallbank, Jennifer E. Dumaine, Amandine Guérin, Ian S. Cohn, Keenan M. O’Dea, Ryan D. Pardy, Maria I. Merolle, Lindsey A. Shallberg, Emma N. Hunter, Jessica H. Byerly, Eleanor J. Smith, Gracyn Y. Buenconsejo, Briana I. McLeod, David A. Christian, Boris Striepen, Christopher A. Hunter

**Affiliations:** Department of Pathobiology, School of Veterinary Medicine, University of Pennsylvania, 380 South University Avenue, Philadelphia, PA 19104, United States of America; Cell Press, 50 Hampshire St, Cambridge, MA 02139, United States of America

## Abstract

*Cryptosporidium* causes debilitating diarrheal disease in patients with primary and acquired defects in T cell function. However, it has been a challenge to understand how this infection generates T cell responses and how they mediate parasite control. Here, *Cryptosporidium* was engineered to express a parasite effector protein (MEDLE-2) that contains the MHC-I restricted SIINFEKL epitope which is recognized by TCR transgenic OT-I CD8^+^ T cells. These modified parasites induced expansion of endogenous SIINFEKL-specific and OT-I CD8^+^ T cells that were a source of IFN-γ that could restrict growth of *Cryptosporidium*. This T cell response was dependent on the translocation of the effector and similar results were observed with another secreted parasite effector (ROP1). Although infection and these translocated effector proteins are restricted to intestinal epithelial cells (IEC), type I dendritic cells (cDC1) were required to generate CD8^+^ T cell responses to these model antigens. These data sets highlight *Cryptosporidium* effectors as targets of the immune system and suggest that crosstalk between enterocytes and cDC1s is crucial for CD8^+^ T cell responses to *Cryptosporidium*.

## Introduction

*Cryptosporidium spp*. are intracellular, yet extracytoplasmic, apicomplexan parasites that infect intestinal epithelial cells (IECs)^1^. These organisms are a common cause of severe diarrheal disease in children^2^ and *Cryptosporidium* is an opportunistic pathogen in patients with primary or acquired defects in T cell function – such as those with HIV or patients treated with immunosuppressive drugs to prevent transplant rejection^3–5^. These clinical examples highlight the importance of T cells in resistance to *Cryptosporidium*; further exemplified by murine models in which CD4^+^ and CD8^+^ T cells and their production of IFN-γ are crucial for parasite control^6–11^. In addition, there are numerous dendritic cell (DC) subsets present in the intestine that may be involved in this response and there is good evidence that type 1 conventional DCs (cDC1) are important for resistance to *Cryptosporidium*^9,12–14^. This DC subset appears to be an important source of IL-12 that stimulates innate and adaptive sources of IFN-γ^7–10,15–17^ and are required for the generation of *Cryptosporidium*-specific CD4^+^ T cell responses^9^.

The lack of reagents to readily study the T cell response to *Cryptosporidium* has limited the ability to dissect the events that lead to the development of long-term resistance to this organism. For example, because *Cryptosporidium* only infects enterocytes and does not readily breach the intestine, it is unclear what factors govern how DCs might encounter and acquire parasite-derived antigens to prime T cell responses. In addition, the “activated but resting” status of intestinal T cells in the intestine makes it difficult to utilize conventional markers of antigen experience (such as levels of CD11a, CD69, or CD44) as surrogates to distinguish *Cryptosporidium*-specific T cell populations versus those specific for other microbial or environmental antigens present in the intestine^18–20^. One solution to be able to reliably identify T cells that are responsive to microbially-derived peptides is to engineer pathogens to express well characterized model antigens. These genetically modified organisms can then be combined with T cell receptor (TCR) transgenic T cells (such as OT-I T cells specific for the SIINFEKL epitope derived from ovalbumin) or MHC-tetramers loaded with relevant peptides to allow the identification of endogenous T cells that have encountered the model antigen. This approach has been utilized for a range of pathogens that include *Listeria monocytogenes, Plasmodium spp., Trypanosoma cruzi*, and *Toxoplasma gondii* to provide tractable systems to understand how T cell responses are generated and function during these infections^21–24^.

For *Cryptosporidium*, the recent development of transgenesis allowed the identification of *Cryptosporidium* effector proteins, some of which were found to be translocated into host cells^25–28^. Thus, MEDLE-2 and ROP1 are secreted into infected cells but with different kinetics and localize to different regions of the infected cell. To determine if the adaptive immune system could respond to these effectors, *Cryptosporidium parvum* (*Cp*) parasites were engineered to express different forms of these molecules that contained a C-terminal SIINFEKL peptide derived from ovalbumin (OVA) and a hemagglutinin (HA) tag to track their patterns of expression. The MEDLE-2 variant (M2-OVA) induced the activation and expansion of OT-I CD8^+^ T cells as well as endogenous SIINFEKL-specific CD8^+^ T cells. Analysis of these parasite-induced CD8^+^ T cells highlighted phenotypic changes associated with exposure to antigen and tissue occupancy during cryptosporidiosis while the ability of these OT-I T cells to produce IFN-γ at the site of infection contribute to parasite control. However, this antigen needed to be secreted into the host IEC to induce a CD8^+^ T cell responses and in mice that lacked cDC1s, there was a major defect in the ability to induce an OT-I T cell response. Thus, the generation of transgenic parasites expressing a model antigen highlight *Cryptosporidium* effectors as targets of the immune system and suggest that crosstalk between enterocytes and cDC1s is crucial for CD8^+^ T cell responses to *Cryptosporidium*.

## Results

### The use of a model antigen to generate parasite-specific CD8+ T cell responses

To study the CD8^+^ T cell response to *Cryptosporidium,* parasites were engineered to express the SIINFEKL epitope and an HA epitope tag as translational fusions to the C-terminus of the secreted effector protein MEDLE-2 (M2)^25^ in the endogenous locus (Fig. 1A, see materials and methods). When HCT-8 cells were infected with these transgenic parasites then co-stained for HA and *Vicia villosa lectin* (VVL – binds glycans and used to stain *Cryptosporidium*^29^), imaging revealed that M2-OVA was translocated to the cytosol of the infected IECs but was not detected in uninfected cells (Fig. 1B). Previous studies have shown that in IFN-γ^-/-^ mice the high parasite burdens facilitate the detection of immune responses to *Cryptosporidium*^8,17^. Therefore, in initial studies to determine whether M2-OVA parasites induce a SIINFEKL-specific CD8^+^ T cell response, congenically disparate CD45.1.2^+^ OT-I T cells were transferred into CD45.2^+^ IFN-γ^-/-^ mice before infection with WT parasites or M2-OVA *Cp*. Analysis of the intraepithelial lymphocyte (IEL) compartment of the ileum at 10 days post infection (dpi) revealed that OT-I T cells failed to expand in naïve mice (data not shown) or mice infected with WT Cp (Fig. 1C). In contrast, infection with M2-OVA parasites generated a robust OT-I response, that at the peak of infection ranged from ∼5-50% of total CD8^+^ T cells across multiple experiments (Fig. 1C). The use of MHC-I tetramers (SIINFEKL:K^b+^) could also detect the presence of an endogenous SIINFEKL-specific CD8^+^ T cell response in the IEL (Fig. 1D). At this time point, a low frequency of parasite-specific CD8^+^ T cells (SIINFEKL:K^b+^ and OT-I T cells) could also be detected in the mesenteric lymph node (mLN) which drain the ileal compartment (Fig. 1D). Thus, incorporation of SIINFEKL with the secreted *Cryptosporidium* effector protein MEDLE-2 results in a low but detectable parasite-induced CD8^+^ T cell responses in the mLN but which was most prominent at the site of parasite replication in the ileum.

**Fig. 1.**
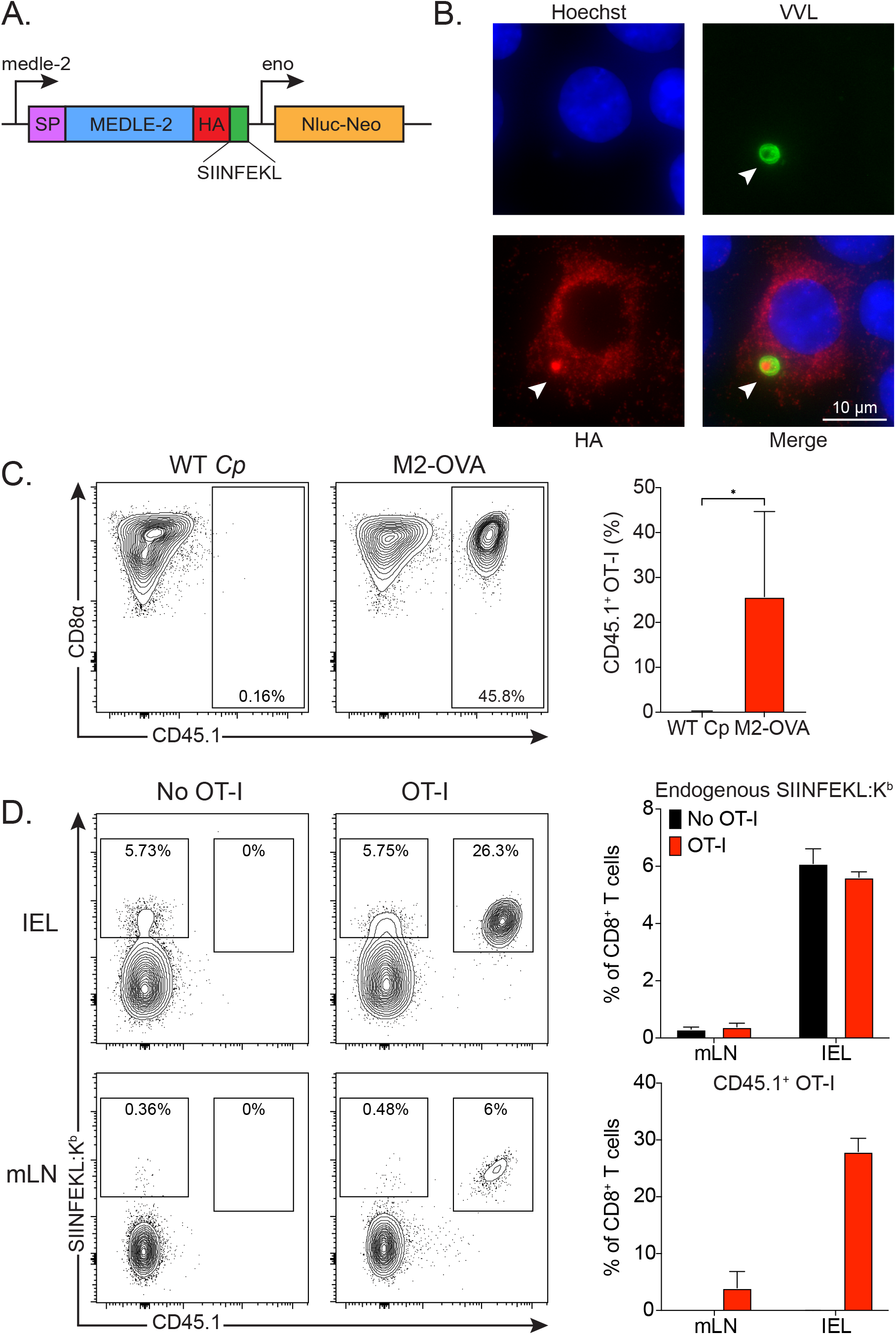
The use of a model antigen to generate parasite-specific CD8^+^ T cell responses. **A.** Genetic construct of transgenic M2-OVA *Cryptosporidium* parasites with the neomycin (Neo) selection marker and the nanoluciferase (Nluc) reporter to monitor parasite burden. **B.** HCT-8 cells were infected for 9 hours then stained for nuclear dye, Hoechst (blue), glycans, *Vicia villosa* lectin conjugated to FITC (VVL) (green), and HA (red). A white arrow points to the parasite within the cell in addition to the HA staining in the parasite. **C.** IFN-γ^-/-^ mice received 10^4^ OT-I cells and were infected with 10^4^ WT Cp or M2-OVA parasites and IEL was harvested at 10 dpi for flow cytometry. Representative flow plots show OT-I cells in the IEL, gated on Singlets, Live, CD45.2^+^, CD19^-^, NK1.1^-^, CD3^+^, CD4^-^, CD8α^+^, SIINFEKL:K^b+^, CD45.1^+^. Summary bar graph showing means of n=2-4 mice/group from 2-6 experiments. **D.** IFN-γ^-/-^ mice received none or 5×10^4^ OT-I then were infected with 10^4^ M2-OVA and mLN and IEL were harvested at 10 dpi for flow cytometry. Representative flow plots show SIINFEKL:K^b+^ cells gated on Singlets, Live, CD19^-^, NK1.1^-^, B220^-^, CD3^+^, CD4^-^, CD8α^+^, SIINFEKL:K^b+^, CD45.1^+^. Summary bar graph of one representative experiment showing means of endogenous SIINFEKL:K^b+^ and OT-I CD8^+^ T cell frequency in mLN and IEL. N=2-4 mice/group from 2 experiments. Statistical significance was determined in **C** by Student’s t test with Welch’s correction * p≤0.05.

### Kinetics and location of *Cryptosporidium*-specific CD8^+^ T cell responses

To better understand where and how frequently these *Cryptosporidium*-specific CD8^+^ T cells encounter their cognate antigen, the M2-OVA parasites were used in combination with OT-I cells that express a Nur77-GFP reporter. Here, expression of Nur77-GFP is used to identify CD8^+^ T cells that have experienced recent (12-24 hours) TCR stimulation, where the geometric mean fluorescence intensity (gMFI) of this GFP signal reflects the strength of TCR engagement^30,31^. IFN-γ^-/-^ mice that received Nur77-GFP OT-I cells were infected with M2-OVA parasites, and on 6, 10, 14, and 18 dpi, the ileal IEL compartment, ileal-draining mLN, and ileal Peyer’s Patches (PP) were harvested. At 6 dpi, few OT-I cells were present in the mLN, and OT-I cells were not readily detected in the IEL or PP (Fig. 2A,B;Supp. Fig. 1A). However, by 10 dpi OT-I cells were present in high frequency in the IEL compartment, their frequency was further increased at 14 dpi, and contraction was observed at 18 dpi (Fig. 2A,B). These kinetics are consistent with the peak of parasite burden at 10 dpi and its rapid decline by 14 dpi (Fig. 2C). To complement the flow cytometry findings, multiphoton imaging approaches were used to visualize OT-I cells in the intestine. IFN-γ^-/-^ mice received 10^6^ GFP^+^ OT-I cells then were infected with 5×10^4^ M2-OVA that also expressed tdTomato to ensure both T cells and parasites could be visualized. Between 5-7 dpi mice were anesthetized and the lumen of an ileal section was exposed to a multiphoton microscope and live imaging revealed that parasite-specific OT-I cells were readily detected in the villi of the ileum near infected cells (Fig. 2D).

**Fig. 2.**
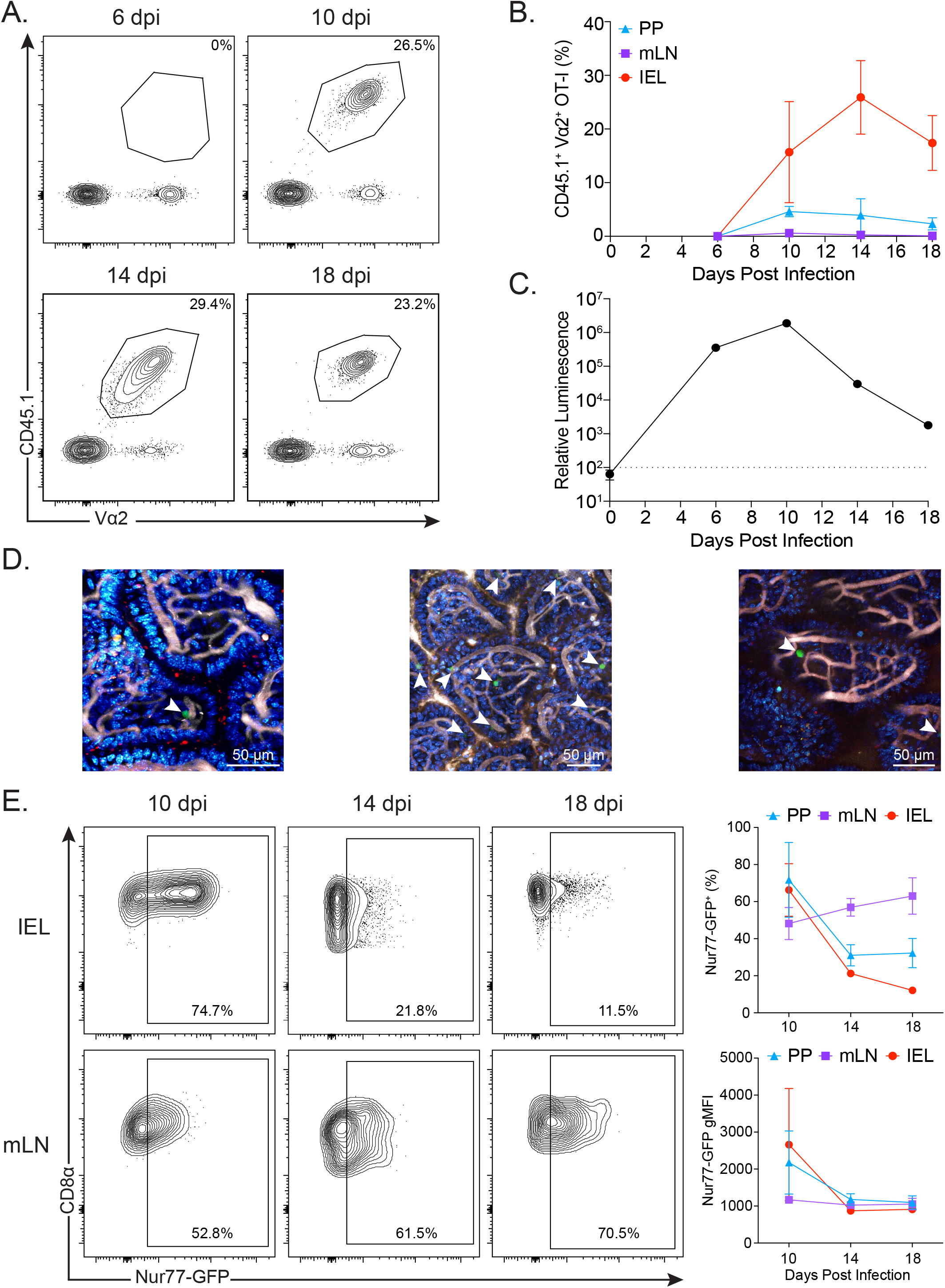
Kinetics and location of *Cryptosporidium*-specific CD8^+^ T cell responses. IFN-γ^-/-^ mice received 10^4^ OT-I cells, were infected with 10^4^ M2-OVA, then PP, mLN, and IEL were harvested at 6, 10, 14, and 18 dpi for flow cytometry. **A.** Representative flow plots display OT-I cells in IEL gated on Singlets, Live, CD19^-^, NK1.1^-^, CD90.2^+^, CD4^-^, CD8α^+^, CD45.1^+^, Vα2^+^. **B.** Summary bar graph of one representative experiment showing percentages of OT-I cells over time. N=2-4 mice/group for 2-3 independent experiments per timepoint. **C.** Parasite burden (relative luminescence) over time measured by nanoluciferase assay. **D.** IFN-γ^-/-^ mice received 10^6^ GFP^+^ OT-I cells (green) and were infected with 5×10^4^ M2-OVA parasites that expressed tdTomato (red). Between 5-7 dpi mice were anesthetized and injected with Hoechst (blue) and Qtracker vascular label (white), then the lumen of an ileal section was exposed to a multiphoton microscope for live imaging. 3 representative still images are shown from one experiment. N=3-4 mice/group for 2 independent experiments. White arrows point to GFP^+^ OT-I cells in the villi of the ileum. **E.** Representative flow plots of Nur77-GFP OT-I cells in IEL and mLN, gated similarly to **A**. Summary bar graph of one representative experiment showing percentages and gMFI of Nur77-GFP^+^ cells. N=2-4 mice/group for 2-3 independent experiments per timepoint.

When the gut-associated secondary lymphoid organs (SLO) were examined 10-18 dpi, a small number of these cells were present in the mLN and PP (Fig. 2A,B;Supp. Fig. 1A). In all three compartments examined a subset of these OT-I cells were Nur77-GFP^+^ (Fig 2E;Supp. Fig. 1B). While the rise and fall of Nur77-GFP expression in the IEL compartment and PP were consistent with parasite burden (Fig. 2E;Supp. Fig. 1B), in the mLN even though there were only low numbers of OT-I T cells present, Nur77-GFP expression appeared to remain elevated between 10 and 18 dpi (Fig. 2E). These data indicate that the highest number of OT-I cells receiving TCR stimulation occur in the IEL compartment at the peak of infection, while the draining LN is a site where OT-I cells receive TCR stimulation over the course of infection.

### *Cryptosporidium*-specific CD8^+^ T cells express markers associated with priming and trafficking to the intestine

Next, this model antigen system was utilized to determine if these T cell populations present in the mLN and IEL could be used to distinguish functional states of activation associated with tissue-specific functions. OT-I cells were transferred into IFN-γ^-/-^ mice and the IEL compartment, mLN, PP, and spleen were harvested at 10 dpi. High-parameter flow cytometry was utilized to assess OT-I cell markers of tissue residency (CD69, PD-1), mucosal association (GzmB, LPAM-1, IL-21R), activation/proliferation (Ki-67, KLRG1, TCF-1), and type 1 immunity (CXCR3, Tbet). The UMAP projection of the aggregate of OT-I cells from all tissues (IEL, PP, mLN, and spleen) illustrates the heterogeneity of the OT-I T cells (Fig. 3A). Projections from each tissue reveals that the OT-I cells present in the mLN and spleen cluster separately from the IEL, while the PP OT-I cells display features of both (Fig. 3A). X-Shift clustering analysis of the aggregated OT-I cells independent of tissue localization identified 6 distinct clusters (Fig. 3B). A heatmap showing the most differentially expressed proteins between these clusters revealed that OT-I cells in clusters 1, 2, 3, and 6 expressed PD-1, and OT-I cells in clusters 2, 3, 4, and 5 expressed LPAM-1 (an integrin that mediates T cell homing to the intestine). While cluster 5 was determined based on LPAM-1 alone, clusters 2, 3, and 4 had additional proteins that defined them. Cluster 2 was based on Ki-67 and PD-1 expression, cluster 3 was determined by PD-1 and TCF-1 expression, and cluster 4 was defined by KLRG1 expression. The flow cytometry data shown in Fig. 3D-I illustrate these different expression profiles. Thus, OT-I cells in the PP and IEL expressed the highest levels of tissue residency (CD69, Fig. 3D), while those in the spleen and mLN expressed significantly higher LPAM-1 (Fig. 3E). In addition, OT-I cells in the mLN and PP have higher levels of replication (Ki-67+, Fig. 3F) compared to cells at the site of infection. Further, OT-I cells in the spleen and mLN have higher KLRG1 expression than OT-I cells in the IEL (Fig. 3G), consistent with recent activation and absence of KLRG1 expression in IEL as previously described^32^. Finally, OT-I cells in all tissues express Tbet and CXCR3, though levels are higher in lymphoid tissues than in the intestine (Fig. 3H, I). Thus, OT-I CD8^+^ T cell responses induced by *Cryptosporidium* have unique tissue-specific markers associated with T cell priming and expansion in the mLN, and trafficking through the periphery before entry and accumulation in the IEL compartment.

**Fig. 3.**
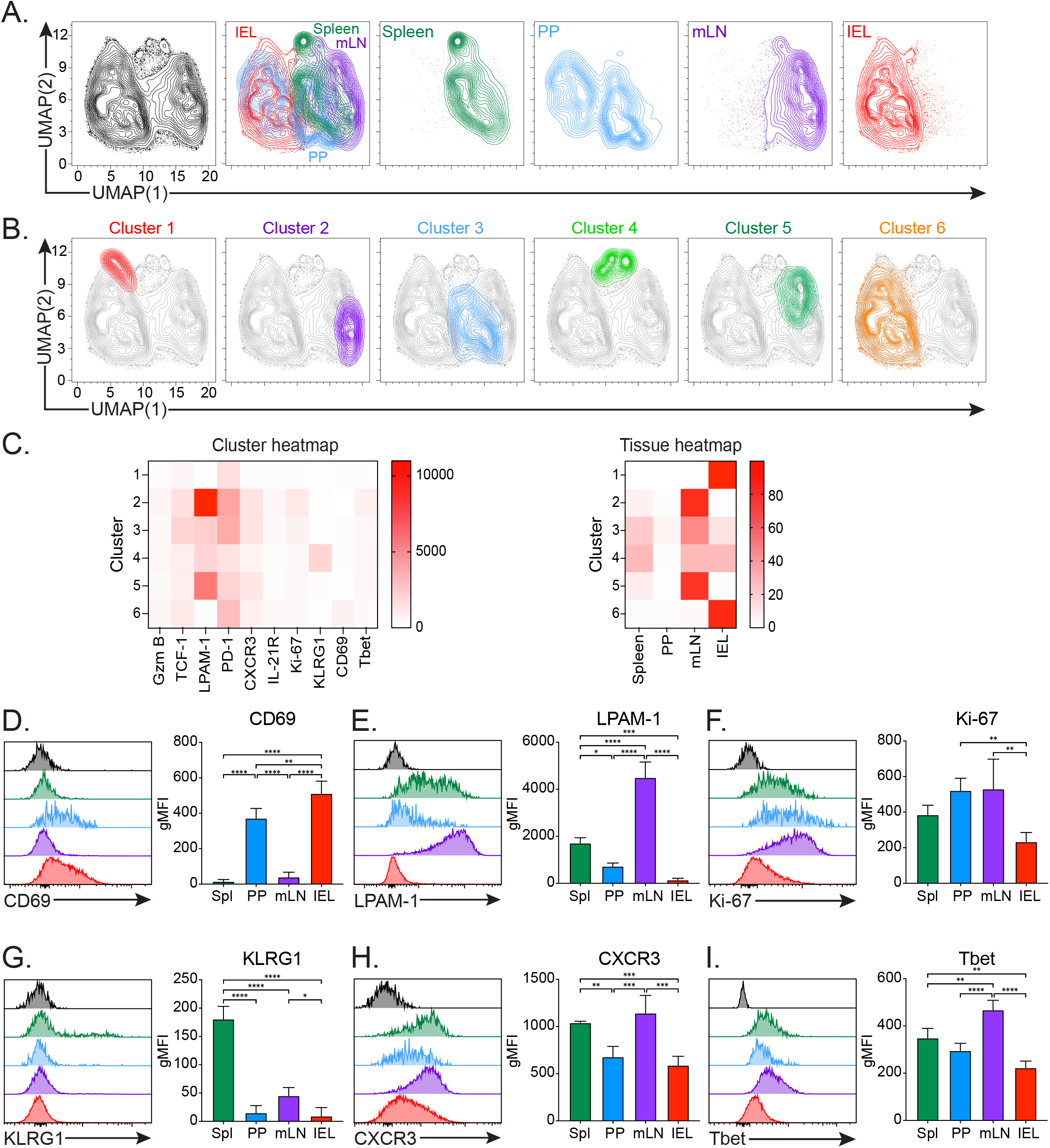
*Cryptosporidium*-specific CD8^+^ T cells express markers associated with priming and trafficking to the intestine. IFN-γ^-/-^ mice received 10^4^ OT-I cells, were infected with M2-OVA, and then spleen, PP, mLN, and IEL were harvested at 10 dpi for flow cytometry. **A.** UMAP based on expression of markers of OT-I cells from all tissues or displaying OT-I cells per tissue. **B.** X-shift cluster analysis of all OT-I cells in UMAP. **C.** Cluster heatmap from X-shift analysis showing expression intensity of markers on all OT-I cells, and a heatmap showing which clusters OT-I cells from each tissue resemble. **D-I**. Representative flow plots of markers on OT-I cells in each tissue, as well as a summary bar graph of the gMFI from one experiment. N=3 mice/experiment for 2 independent experiments, except for spleen which has only been performed once. Statistical differences for **D-I** were determined based on two-way ANOVA and multiple comparisons. * p≤0.05, ** p≤0.01, *** p≤0.001, **** p≤0.0001.

### *Cryptosporidium*-specific CD8^+^ T cells produce IFN-γ during infection

Next, mice that express Thy1.1 transcripts bi-cistronically under the control of the IFN-γ promoter were utilized to detect those T cells that produced IFN-γ during infection^33^. In these experiments, IFN-γ-Thy1.1 reporter mice were treated with αIFN-γ to boost parasite burden and infected with M2-OVA, and the endogenous CD8^+^ T cells were assessed for Thy1.1 expression at 10 dpi. In the mLN and PP of uninfected and infected mice, CD8^+^ T cell Thy1.1 expression was low (Fig. 4A;Supp. Fig. 2). In the IEL compartment of naïve mice, less than 1% of CD8^+^ T cells expressed Thy1.1 (Fig. 4A). At 10 dpi, CD8^+^ T cells in the IEL showed a ten-fold increase in Thy1.1 expression, and ∼5-30% of Thy1.1^+^ cells stained for SIINFEKL:K^b^ tetramer (Fig. 4A). To determine if the ability of the parasite-specific CD8^+^ T cells to produce IFN-γ would enhance protection against *Cryptosporidium*, 10^6^ OT-I cells were transferred into IFN-γ^-/-^ mice, which were then infected with M2-OVA parasites. In mice that did not receive OT-I cells parasite burden peaked at 7 dpi, declined by 10 and 12 dpi, but remained at low levels at later time points (Fig. 4B). Based on area under the curve analysis, the transfer of 10^6^ OT-I cells consistently displayed ∼75% reduced parasite burdens, but was not sufficient to promote parasite clearance (Fig. 4B). To determine whether the protection provided by OT-I cells was dependent on IFN-γ, IFN-γ^-/-^ mice received 10^6^ OT-I cells, were treated with either isotype or αIFN-γ antibodies, and parasite burden was assessed. Based on area under the curve analysis for 8-18 dpi across multiple experiments, mice that received OT-I cells and αIFNγ antibody had ∼175% increased parasite burdens compared to mice that received OT-I cells and isotype antibody (Fig. 4C). These data indicate that the ability of OT-I T cells to respond to *Cryptosporidium*-derived antigens and produce IFN-γ can contribute to parasite control.

**Fig. 4.**
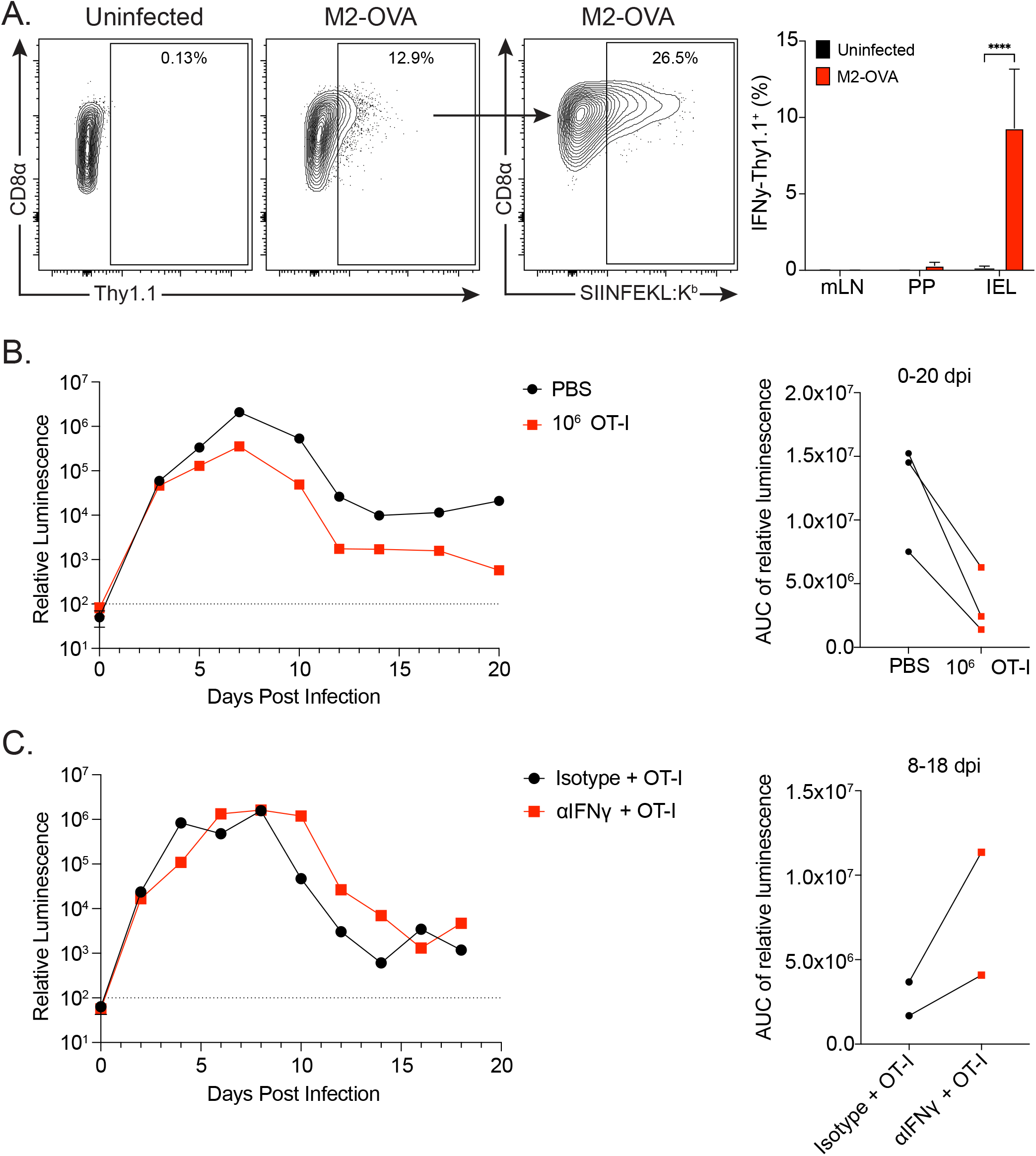
*Cryptosporidium*-specific CD8^+^ T cells produce IFN-γ during infection. **A.** IFN-γ-Thy1.1 reporter mice were left untreated and uninfected or treated with 1 mg/mouse αIFN-γ 2 days prior to infection and 2 days post infection with 10^4^ M2-OVA. mLN, PP, and IEL were harvested 10 dpi for flow cytometry. Representative flow plots show Thy1.1 cells in IEL (left), gated on Singlets, Live, CD19^-^, NK1.1^-^, CD90.2^+^, CD4^-^, CD8α^+^, Thy1.1^+^. Representative flow plot shows Thy1.1^+^ cells from infected mice gated further down on SIINFEKL:K^b^ (right). Summary bar graphs from one experiment of percentages of Thy1.1^+^ cells. N=2-4 mice/group for 2 independent experiments. **B.** PBS or 10^6^ OT-I cells were transferred into IFN-γ^-/-^ mice, infected with 10^4^ M2-OVA, and feces were analyzed by nanoluciferase assay for parasite burden (relative luminescence) over time. Area under the curve analysis was performed for each treatment 0-20 dpi for each independent experiment. N=3 mice/group for each experiment and experiments were performed 3 times. **C.** IFN-γ^-/-^ mice were treated with 1 mg/mouse isotype IgG1 anti-horseradish peroxidase antibody or αIFN-γ 1 day prior to infection as well as 2, 5, and 8 days post infection that were infected with 10^4^ M2-OVA. Feces was analyzed by nanoluciferase assay for parasite burden (relative luminescence) over time. Area under the curve analysis was performed for each treatment for each independent experiment from 8-18 dpi. N=3-4 mice/group for each experiment and were performed 2 times. Statistical differences in **A** were determined based on two-way ANOVA and multiple comparisons. * p≤0.05, ** p≤0.01, *** p≤0.001, **** p≤0.0001.

### Secretion of *Cryptosporidium* antigens is required to induce CD8^+^ T cell responses

In the studies described above, the M2-OVA construct results in SIINFEKL expression in the parasite and a portion of this is translocated to the host cytosol. To determine whether antigen localization influences the ability to generate CD8^+^ T cell responses, several additional transgenic parasite lines were generated. One was a variant of the M2-OVA construct which lacked the signal peptide required for co-translational insertion into the endoplasmic reticulum and secretion^25^ (NS-M2-OVA, Fig. 5A). In addition, SIINFEKL was conjugated to ROP1 (ROP1-OVA), an effector protein that is stored in the rhoptry organelle, and then injected into the host cell only during invasion^26^ (Fig. 5A). A comparison of localization of these constructs in HCT-8 cells highlighted that M2-OVA resulted in high levels of antigen distributed through the cytosol of the infected cell, whereas the NS-M2-OVA parasites showed that the tagged antigen remained localized within the parasite (Fig. 5B). In contrast to the M2-OVA construct, when HCT-8 cells were infected with ROP1-OVA parasites, low levels of tagged protein were exported to the host cell cytoplasm with a punctate expression pattern associated with the site of parasite invasion and at the cell periphery (Fig. 5B), likely due to its association with the cortical cytoskeleton of the host cell^26^.

**Fig. 5.**
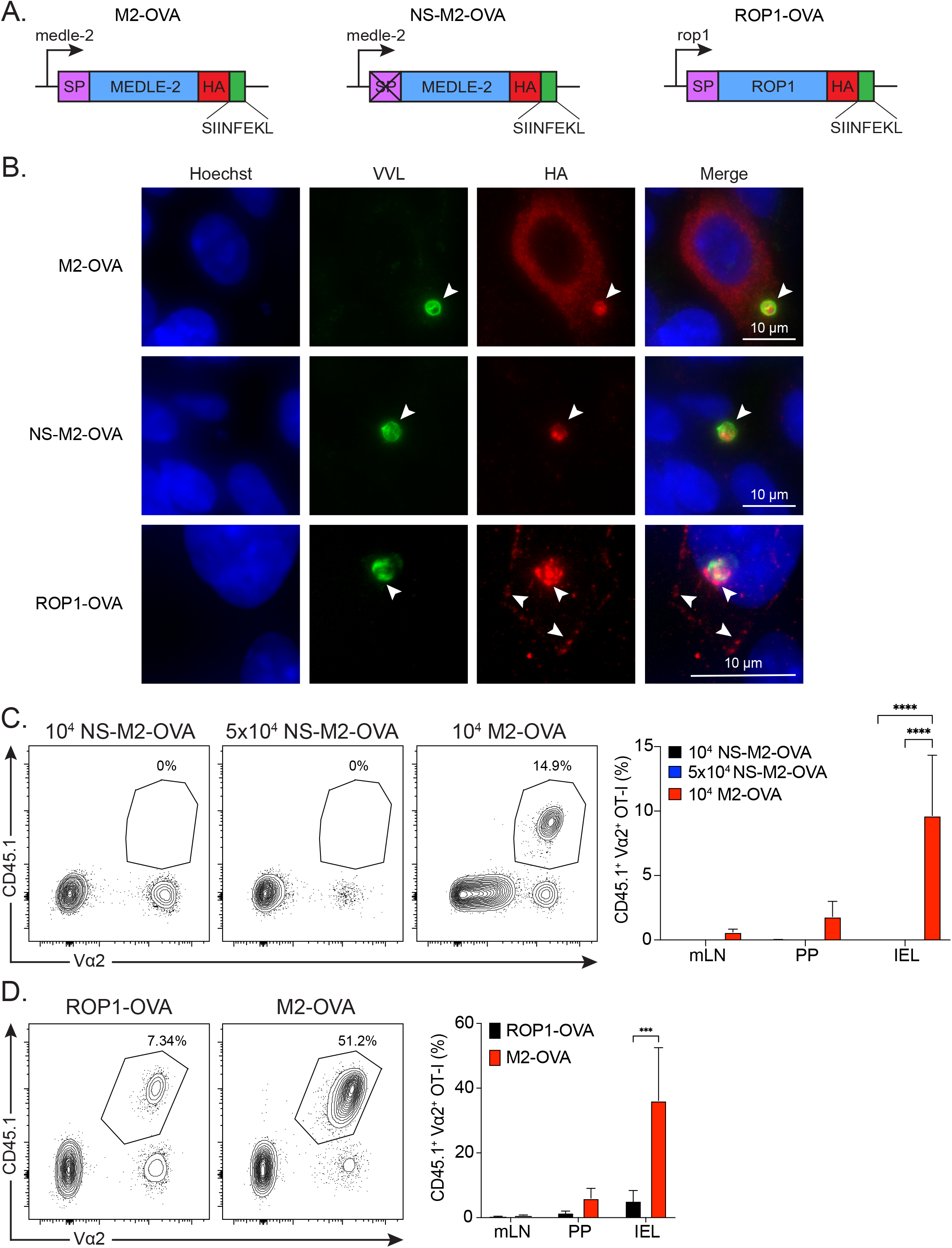
Secretion of *Cryptosporidium* antigens is required to induce CD8^+^ T cell responses. **A.** Genetic constructs of transgenic M2-OVA, NS-M2-OVA, and ROP1-OVA *Cryptosporidium* parasites are shown. **B.** HCT-8 cells were infected with M2-OVA or NS-M2-OVA for 9 hours then stained for nuclear dye, Hoechst (blue), glycans, *Vicia villosa* lectin conjugated to FITC (VVL) (green), and HA (red). A white arrow points to the parasite within the cell in addition to the HA staining in the parasite. Additionally, HCT-8 cells were infected with ROP1-OVA for 2 hours then stained for nuclear dye, Hoechst (blue), glycans, VVL (green), and HA (red). White arrows point to the parasite within the cell as well as the HA staining near the site of infection and around the cell periphery. **C.** IFN-γ^-/-^ mice received 10^4^ OT-I cells and were infected with 10^4^-5×10^4^ NS-M2-OVA or M2-OVA parasites and mLN, PP, and IEL were harvested at 10 dpi for flow cytometry. Representative flow plots show OT-I cells in IEL gated on Singlets, Live, CD19^-^, NK1.1^-^, CD90.2^+^, CD4^-^, CD8α^+^, CD45.1^+^, Vα2^+^. Summary bar graph of one representative experiment showing percentages of OT-I cells in each tissue. N=2-3 mice/group for each experiment and has been performed 3 times. The 5×10^4^ NS-M2-OVA infection group was only performed once. **D.** IFN-γ^-/-^ mice received 10^4^ OT-I cells and were infected with 10^4^ ROP1-OVA or M2-OVA parasites and mLN, PP, and IEL were harvested at 10 dpi for flow cytometry. Representative flow plots show OT-I cells in IEL gated similarly to **C**. Summary bar graph of one representative experiment showing percentages of OT-I cells in each tissue. n=3 mice/group from 3 independent experiments. Statistical differences in **C** and **D** were determined based on two-way ANOVA and multiple comparisons. * p≤0.05, ** p≤0.01, *** p≤0.001, **** p≤0.0001.

To examine whether these altered patterns of expression impacted T cell activation, 10^4^ OT-I cells were transferred into IFN-γ^-/-^ mice that were then infected with M2-OVA or NS-M2-OVA parasites. At 10 dpi, M2-OVA induced OT-I expansion in the IEL, mLN, and PP, while infection with NS-M2-OVA did not result in detectable OT-I responses (Fig. 5C;Supp. Fig. 3A). Furthermore, the use of a five-times higher infection dose of the NS-M2-OVA parasites still did not lead to appreciable T cell activation (Fig. 5C;Supp. Fig. 3A). To determine whether the generation of parasite-specific CD8^+^ T cells was a conserved feature of translocated proteins, a direct comparison of the ability of the M2-OVA and the ROP1-OVA parasites to induce expansion of the OT-I T cells was performed. Across multiple experiments, the ROP1-OVA infected mice reliably induced OT-I responses (Fig. 5D;Supp. Fig. 3B). However, in the experiment shown the magnitude of the response appeared to reflect differences in parasite burden for the two strains (Supp. Fig. 3C). Therefore, it is unclear if the differences in magnitude of the OT-I response were due to parasite burden or underlying biology of the two secreted proteins. For example, ROP1 is injected as a single event during invasion while MEDLE-2 is constitutively translocated from 6 hours post infection. Nevertheless, the use of these transgenic parasite lines indicates that translocation of *Cryptosporidium* antigens into the cytosol of the infected cell leads to the priming and expansion of parasite-specific CD8^+^ T cells.

### cDC1s are required for *Cryptosporidium*-specific CD8^+^ T cell responses

Previous studies have indicated that DCs, specifically cDC1s, and IL-12 production are required for CD4^+^ T_h_1 responses and thereby promote resistance to *Cryptosporidium*^9,12–14^. However, it is unclear if cDC1s are necessary for CD8^+^ T cell responses to *Cryptosporidium*. To address this question, OT-I cells were transferred into WT C57BL/6J or *Irf8+32*^-/-^ mice, which lack cDC1s^34^, that were then infected with M2-OVA parasites. As previously reported, *Irf8+32*^-/-^ mice were more susceptible to infection (∼1000% increase compared to WT) and did not resolve this infection (Fig. 6A). Despite this increase in parasite burden, *Irf8+32*^-/-^ mice had a significant defect in the generation of OT-I T cell responses in the IEL, but no statistical differences in the mLN and PP (Fig. 6B; Supp. Fig. 4). Together, these data demonstrate that cDC1s are required for intestinal CD8^+^ T cell responses to *Cryptosporidium*.

**Fig. 6.**
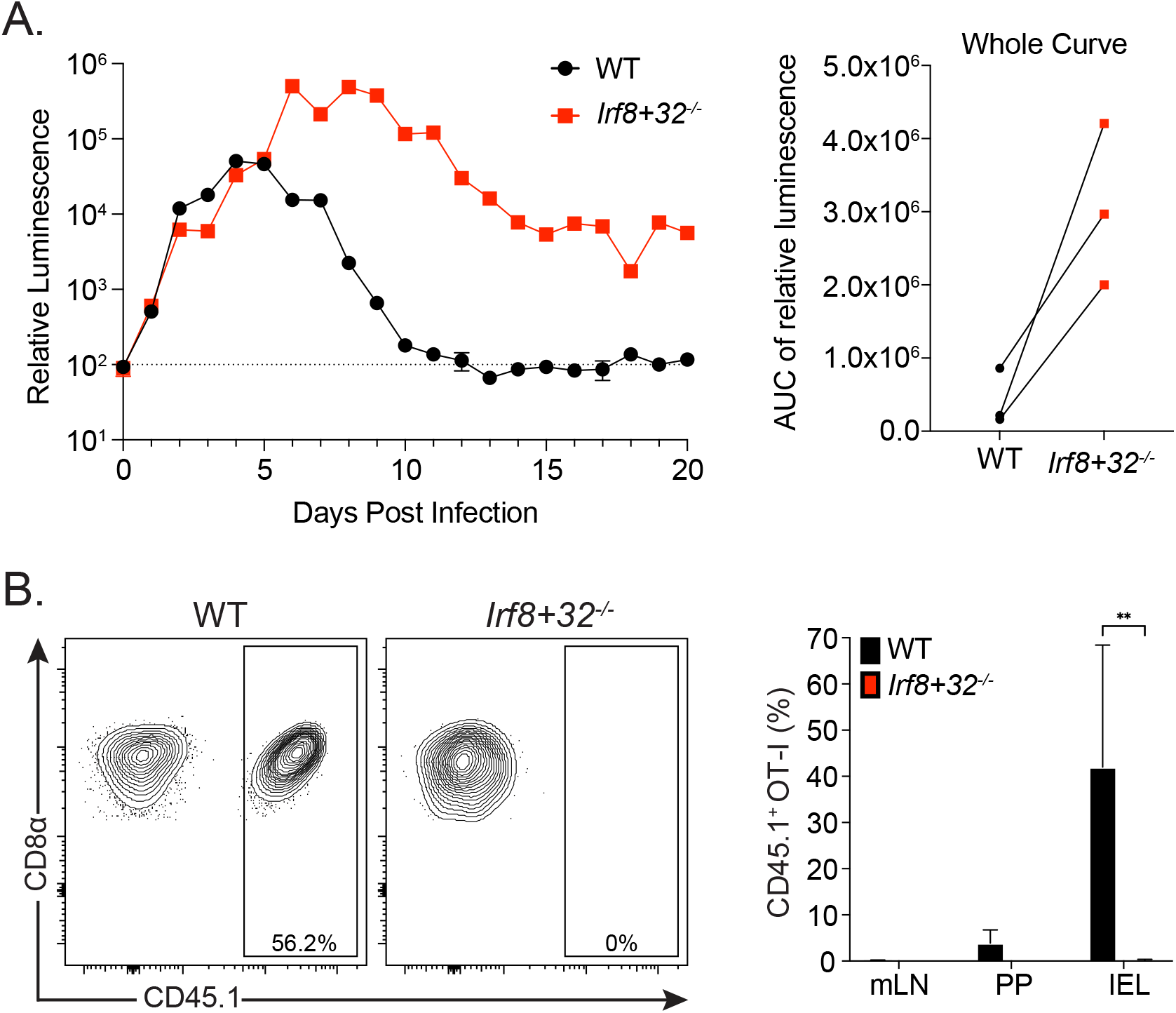
cDC1s are required for *Cryptosporidium*-specific CD8^+^ T cell responses. **A.** WT C57BL/6J or *Irf8+32*^-/-^ mice were infected with 10^4^ M2-OVA parasites and feces analyzed by nanoluciferase assay for parasite burden (relative luminescence) over time. Area under the curve analysis was performed for each treatment for each independent experiment. N=3 mice/group and was performed 3 times. **B.** 10^4^ OT-I cells were transferred to WT or *Irf8+32*^-/-^ mice, infected with 10^4^ M2-OVA, then mLN, PP, and IEL were harvested at 10 dpi for flow cytometry. Representative flow plots show OT-I T cells in IEL gated on Singlets, Live, CD19^-^, NK1.1^-^, CD90.2^+^, CD4^-^, CD8α^+^, CD45.1^+^. Summary bar graph of one representative experiment showing percentages of OT-I T cells in each tissue. N=3 mice/group for each experiment and was performed 2 times.

## Discussion

While it is known that T cells contribute to control of *Cryptosporidium*^6–9,11^, the paucity of tools to readily identify *Cryptosporidium*-specific T cells has hindered the ability to understand the events that influence the development of protective immunity. Recently, TCR sequencing was performed to identify CD4^+^ T cell clones that were specific to *Cryptosporidium* antigens^9^, and a natural *Cryptosporidium* MHC-I restricted epitope was identified that induced CD8^+^ T cell responses during infection^35^. The application of transgenesis to engineer *Cryptosporidium* that express different variants of a model antigen provides a complimentary approach to dissect the basis for resistance to *Cryptosporidium*. Thus, in this experimental system, infection of IFN-γ-Thy1.1 reporter mice established that SIINFEKL-specific CD8^+^ T cells produced IFN-γ while the transfer of OT-I T cells into IFN-γ^-/-^ mice mediated IFN-γ-dependent protection against *Cryptosporidium*. However, this potent CD8^+^ T cell activity directed towards a single peptide was not sufficient for parasite clearance, indicating that additional T cell specificities to other parasite antigens are required for sterile immunity. Indeed, the use of the IFN-γ-Thy1.1 reporters highlighted that infection resulted in an overall increase in CD8^+^ T cell responses, but it was difficult to distinguish whether this was a response to *Cryptosporidium* derived antigens or a secondary consequence of infection-induced inflammation and responses to gut-associated commensals^36,37^. While IFN-γ is perhaps considered the major mediator of resistance to *Cryptosporidium*, there are IFN-γ-independent, T cell-dependent mechanisms of control^8^, which may include the ability of CD8^+^ T cells to lyse infected cells^11^. Activated OT-I cells present in the IEL compartment expressed granzyme B, a key component for cytotoxicity, and the ability to use intravital microscopy should provide the opportunity to visualize whether CD8^+^ T cells can interact directly with and lyse infected cells *in vivo*.

The ability to track the OT-I T cell responses to parasite-derived SIINFEKL provided the opportunity to better understand where and when they encountered cognate antigen and how this would influence T cell function. For example, the use of Nur77-GFP reporter OT-I cells showed that, for the few cells present in the mLN, TCR engagement appeared sustained throughout infection. In contrast, in the IEL compartment this was more dynamic and correlated strongly with parasite burden and the local production of IFN-γ. Further analysis of OT-I cells at 10 dpi indicated that in the IEL these cells displayed a profile of markers consistent with tissue occupancy and exposure to cognate antigen. For example, CD69 expression was high in the IEL but comparatively reduced Tbet and KLRG1 expression in the IEL is reminiscent of tissue resident memory cells (Trm)^32,38–42^. This model system should provide an opportunity to generate *Cryptosporidium*-specific memory CD8^+^ T cell responses and characterize their durability and ability to mediate protection to secondary challenges.

DCs are considered important antigen presenting cells and a source of IL-12 required for resistance to *Cryptosporidium*^12–14^. However, the role of individual DC subsets in the distinct processes of initial priming versus the regulation of effector responses at the local site of infection are uncertain. In other models, cDC1 are predominantly associated with the process of cross-presentation (the ability to sample antigen from the environment) to activate naïve CD8^+^ T cells. In contrast, previous studies with *Cryptosporidium* have highlighted that the ability of cDC1 to produce IL-12 induces CD4^+^ T cell responses dominated by the production of IFN-γ^9^. The data presented here indicate that cDC1s are required for the priming/expansion of *Cryptosporidium*-specific CD8^+^ T cells but do not distinguish the relative contribution of cDC1 antigen presentation and cytokine production to OT-I T cell activation. Since DCs are not infected by *Cryptosporidium*, it is unknown how these antigen presenting cells (APCs) might acquire antigen for processing. One possibility is that parasite secretion of antigen into the IEC cytoplasm is required for DC to acquire antigen for presentation to T cells. Whether this would require active crosstalk between IEC and DC or death of the infected cell for antigen transfer is unclear. Another option includes the ability of goblet cells to acquire luminal antigen and then use goblet cell associated antigen passages (GAPS) to transfer it to DCs in the lamina propria^43^. Similarly, CX3CR1^+^ APCs are specialized to sample intestinal lumen antigen, then translocate antigen to CX3CR1^-^ DCs to activate T cell responses^44–46^. These prior studies on mechanisms of antigen acquisition have been performed with either soluble antigens or enteric pathogens, such as *Salmonella,* that breach the intestinal barrier and may not be relevant to *Cryptosporidium*. However, a recent study that utilized mice in which enterocytes express an ovalbumin-flagellin fusion protein found that activation of the inflammasome to induce cell death promoted cross presentation of SIINFEKL by DCs to prime CD8^+^ T cells^47^. It is relevant to note that the NLRP6 inflammasome has an important role in innate resistance to *Cryptosporidium*^48^, but whether this contributes to antigen presentation during *Cryptosporidium* infection is uncertain. Nevertheless, the experimental approaches described here may help to define how DC can access *Cryptosporidium*-derived antigens required to generate protective T cell responses.

Previous studies indicated that the *Cryptosporidium* export machinery does not readily process rigorously folded proteins for translocation into the host cell, whereas MEDLE-2 is disordered and efficiently exported^25^. Other apicomplexan parasites, such as *Plasmodium* and *Toxoplasma,* have similar export and translocation machinery^49,50^ and *Plasmodium* parasites have been engineered to express SIINFEKL peptide that is secreted into the hepatocyte cytosol^51^. In contrast, *Toxoplasma* parasites can secrete larger OVA fragments into the parasitophorous vacuole^52^, but the ability to transport beyond the vacuole may be restricted to disordered cargo^53,54^. Using transgenes that can be specifically engineered provides control over when the antigen is expressed, and to which subcellular compartment it is localized. For other pathogens, the ability to target model antigens to different compartments (intracellular, surface, or secreted) impacts the efficiency with which these are recognized^22,24,52,55^. In several experimental systems, secreted model antigens are preferentially recognized^22,24,52,55^ and this is reflected in instances where secreted pathogen-derived molecules generate strong CD4^+^ and CD8^+^ responses^52,55,56^. Similarly, when *Cryptosporidium* antigen was not secreted into the host cytoplasm, there was no OT-I T cell response. This observation suggests that exported or secreted antigens from *Cryptosporidium* may be the dominant MHC-I restricted antigens that are presented by DCs. A practical implication of this work is that secreted parasite antigens may represent ideal candidates for vaccine efforts designed to generate cell-mediated immunity to *Cryptosporidium*. In other words, pathogen-derived effectors that interface with host cellular machinery represent a point of weakness that can be exploited by the host immune system. However, this can also manifest as a selective pressure on the pathogen that is reflected in the highly polymorphic nature of proteins associated with the secretory organelles of *Cryptosporidium*^28^.

## Materials and Methods

### Mice

C57BL/6 (stock no: 000664), Nur77-GFP reporter mice, (stock no: 016617), CD45.1 C57BL/6 mice (stock no: 002014), *Irf8+32*^-/-^ mice (stock no: 032744), UBC-GFP mice (stock no: 004353), OT-I mice (stock no:003831), and IFN-γ^-/-^ (stock no: 002287) were purchased from Jackson Laboratories and then maintained in-house. IFN-γ-Thy1.1 knock-in reporter mice were provided by Dr. Phillip Scott but originated in the laboratory of Dr. Casey Weaver^33^. Mice used in this study were males or females ranging from 5 to 12 weeks, and all mice were age and sex matched within individual experiments. All protocols for animal care were approved by the Institutional Animal Care and Use Committee of the University of Pennsylvania (protocol #805405 and #806292).

### Plasmid Construction

To see the full list of primers used for plasmid construction see Supp. Table 1. To generate the M2-OVA and ROP1-OVA lines, the SIINFEKL fragment was inserted into p3XHA_nLuc-Neo^26,27^ by Gibson assembly. Repair templates were amplified from the resulting plasmid with primers containing 30 bp overhangs either side of a Cas9 guide-induced double strand break at the C-terminus of either M2 (Cgd5_4590) or ROP1 (Cgd3_1770). Guide sequences were introduced into a Cas9 expressing vector at a BbsI restriction site as previously described^27^. To generate the M2-OVA tdTomato plasmid, tdTomato was inserted by Gibson assembly into p3XHA-SIINFEKL_nLuc-Neo, linked to nLuc-Neo with a T2A skip peptide. To generate the NS-M2-OVA plasmid the SIINFEKL coding sequence was introduced into pΔSP^25^ that expresses a copy of M2 lacking the 22 amino acid signal peptide coding region. The repair template was amplified with 30 bp overhangs either side of a Cas9 guide-induced double strand break in the thymidine kinase (Cgd5_4440) locus^27^.

### Isolation of transgenic parasites

Transgenic parasites were generated as previously described^57^. Briefly, to excyst parasites, oocysts were bleached, washed in PBS, and incubated in sodium taurodeoxycholate. Then sporozoites were resuspended in transfection buffer supplemented with 100 μg DNA (comprising 50 μg of Cas9/gRNA plasmid and 50 μg of repair template) and electroporated using an Amaxa 4D nucleofector (Lonza, Basel, Switzerland). Parasites carrying a stable transgene were selected using paromomycin added to the drinking water of infected mice. Transgenic parasites were propagated by orally infecting IFN-γ^-/-^ mice and oocysts were purified from their feces using sucrose flotation followed by a cesium chloride gradient, as previously described^8^.

### Mouse infection and measurement of parasite burden

Mice were infected with 10^4^-5×10^4^ oocysts by oral gavage. Infected WT C57BL/6J mice were treated with 1 mg αIFN-γ antibody or 1 mg rat IgG1 isotype control, anti-horseradish peroxidase (BioXCell) prior to and after infection on various days, depending on the experiment, which is explained in the figure legends. To quantify fecal oocyst shedding, 20mg of pooled cage feces was suspended in 1mL lysis buffer. Samples were shaken with glass beads for 5 min, then combined in a 1:1 ratio with Nano-Glo® Luciferase solution (Promega, Ref N1150). A Promega GloMax plate reader was used to measure luminescence. Pooled samples were used because previous studies have demonstrated that mice within each cage are equally infected^58^.

### T cell transfers

For T cell transfers, OT-I mice were interbred with CD45.1/Nur77-GFP reporter mice. To isolate OT-I CD8^+^ T cells, lymph nodes and spleens were harvested and leukocytes were obtained by dissociation over a 70 um filter. Red blood cells were lysed by incubation for 5 min at room temperature in 1 mL of lysis buffer (0.864% ammonium chloride (Sigma-Aldrich) diluted in sterile deionized H_2_O), and then washed with complete RPMI (RPMI, 10% fetal calf serum, 0.1% beta-2-mercaptoethanol, 1% non-essential amino acids, 1% sodium pyruvate, and 1% pen-strep). OT-I T cells were enriched by magnetic activated cell sorting (MACS) using the CD8a+ T Cell Isolation Kit (Miltenyi Biotec) or using the EasySep^TM^ Mouse CD8+ T Cell Isolation Kit (Stem Cell Technologies). OT-I purity was verified (∼80-95%) using flow cytometry for TCR Vα2 and Vβ5.1 expression. 10^4^-10^6^ OT-I T cells were transferred by intraperitoneal injection into recipient mice.

### Flow cytometry

Single-cell suspensions were prepared from intestinal sections by shaking diced tissue at 37°C for 20-30 min in Hank’s Balanced Salt Solution with 5 mM EDTA and 1 mM DTT. Cell pellets were then passed through 70 um and 40 um filters. Ileal draining mesenteric lymph nodes and Peyer’s patches from the ileum as well as spleens were harvested and dissociated through 70 um filters, then washed with cRPMI. Cells were washed in FACS buffer (1x PBS, 0.2% bovine serum antigen, 1 mM EDTA), and incubated in Fc block (99.5% FACS Buffer, 0.5% normal rat IgG, 1 µg/ml 2.4G2) at 4°C for 15 min prior to staining. Cells were stained for cell death using either LIVE/DEAD Fixable Aqua Dead Cell marker (Invitrogen) or GhostDye Violet 510 Viability Dye (TONBO Biosciences) in 1x PBS at 4°C for 15 min. Cells were washed after cell death staining and surface antibodies were added and stained at 4°C for 20-30 min. If intracellular staining for transcription factors was performed, cells were fixed using the eBioscience Foxp3 Transcription Factor Fixation/Permeabilization Concentrate and Diluent (ThermoFisher Scientific) for 20 min at 4°C, and then washed with FACS buffer. Cells were then stained for transcription factors in 1x eBioscience Permeabilization Buffer (ThermoFisher Scientific) at 4°C for 30 min. Cells were then washed in FACS buffer prior to acquisition. Cells were stained using the following fluorochrome-conjugated antibodies: CD19 PerCP-Cy5.5 (clone ID3), NK1.1 PerCP-Cy5.5 (clone PK136), CD4 BV650 (clone RM4-5), CD3 BV785 (clone 17A2), CD45.2 BV711 (clone 104), TCR Vb5.1,5.2 APC (clone MR9-4), TCR Va2 PE (clone B20.1), B220 PerCP-Cy5.5 (clone RA3-6B2), CD8a BV650 (53-6.7), CD4 BV711 (clone GK1.5), CD45.1 PE-Cy7 (clone A20), CD45.1 BV711 (clone A20), CD45.2 APC (clone 104), CD90.2 AF700 (clone 30-H12), CD8b FITC (clone YTS156.7.7), CD8a BV711 (clone 53-6.7), CD44 BV785 (clone IM7), CD90.2 BV785 (clone 30-H12), CD4 AF700 (clone RM4-5), PD-1BV605 (clone 29F.1A12), CXCR3 BV650 (clone CXCR3-173), KLRG1 BV711 (clone 2F1), IL-21R APC (clone 4A9), CD45.1 PE-Cy7 (clone A20), IL-21R PE-Cy5 (clone 4A9), CD45.2 PerCP-Cy5.5 (clone 104), CD45.1 BV510 (clone A20), CD11a PE (clone M17/4) from Biolegend; CD90.1 APC-ef780 (clone HIS51), CD8a PE-Cy7 (clone 53-6.7), CD45.1 APC-ef780 (clone A20), CD8a APC-ef780 (clone 53-6.7), TCR Va2 Superbright 780 (clone B20.1), CD45.1 APC (clone A20), Gzm B ef450 (clone NGZB), LPAM-1 PE (clone DATK32), Tbet PE-Cy5 (clone 4B10), CD3e PE-Cy7 (clone 145-2C11), CD8b APC-ef780 (clone eBioH35-17.2), CD4 ef450 (clone GK1.5), Tbet PE-Cy7 (clone eBio4B10), CD45.2 ef450 (clone 104) from eBioscience; CD8a PE-cf594 (clone 53-6.7), CD8a BUV563 (clone 53-6.7), CD19 BUV395 (clone ID3), NK1.1 BUV395 (clone PK136), CD4 BUV805 (clone GK1.5), CD69 BUV737 (clone H1.2F3), CD11a BUV805 (clone 2D7), Ki-67 AF700 (clone B56), EpCAM BUV395 (clone G8.8), CD4 BUV496 (clone GK1.5), CD8a BUV805 (clone 53-6.7), Va2 BV605 (clone B20.1), CD44 PE-Cy5 (clone IM7) from BD Biosciences; TCF1 AF488 (clone C63D9) from Cell Signaling. Endogenous and OT-I responses were measured by H2-K^b^:SIINFEKL tetramer conjugated to PE or APC (NIH Tetramer Core) staining at room temperature for 20-30 min. Data were collected on a FACSCanto, LSRFortessa, or FACSymphony A3 Lite (BD Biosciences) and analyzed with FlowJo v10 software (TreeStar). Cells were downsampled to a consistent number depending on the tissue using Downsample plugin in FlowJo prior to UMAP. UMAP and X-Shift plugins were utilized in FlowJo for dimensionality reduction and unsupervised clustering, and X-Shift was visualized by Cluster Explorer.

### Immunofluorescence imaging

Human ileocecal adenocarcinoma cells (HCT-8) (ATCC) were grown on coverslips in DMEM supplemented with 10% cosmic calf serum (CCS) (Thermo Fisher). 200,000 purified oocysts were excysted and seeded on coverslips when HCT-8 cells were 80% confluent. After excystation at indicated time points post infection, cells were washed with PBS and successively fixed for 10 minutes with 4% paraformaldehyde followed by permeabilization for 10 minutes with 0.1% Triton X-100 (Sigma). Coverslips were blocked with 4% bovine serum albumin (BSA) (Sigma) and antibodies were diluted in 1% BSA solution. Rat monoclonal anti-HA (clone: 3F10; Sigma) was used as primary antibody and goat anti-rat polyclonal Alexa Fluor 594 (Thermo Fisher) as secondary along with *Vicia villosa* lectin conjugated to FITC (Vector Labs). Host and parasite nuclei were stained with Hoechst 33342 (Thermo Fisher). Slides were imaged using a Leica Widefield microscope.

### Multiphoton imaging

GFP-expressing OT-I T cells were isolated from UBC-GFP mice that were interbred with OT-I mice and purified as previously described. IFN-γ^-/-^ mice were injected intraperitoneally with 10^6^ GFP^+^ OT-I cells 24 hours prior to infection with 5×10^4^ *C. parvum* M2-OVA tdTomato parasites. At 5-7 days post infection mice were orally gavaged with 200 µl 2mg/ml loperamide hydrochloride (Sigma Aldrich) to minimize peristaltic movement for imaging. Mice were subsequently anaesthetized by intraperitoneal injection with a xylazine/ketamine cocktail followed by retroorbital injection with 0.5 mg Hoechst 33342 (Invitrogen) and 30 µl Qtracker 655 vascular label (Invitrogen). Mice were maintained under anesthesia at 37°C with vaporized isoflurane. The ileal loop was extracted through a small incision made in the abdomen of the mouse and a c.1 cm section dissected longitudinally to expose the luminal surface. The cut edges were cauterized to minimize blood loss. 20 µg/ml loperamide hydrochloride (Sigma Aldrich) and 20 µg/ml indomethacin (Sigma Aldrich) was applied topically to the luminal intestinal surface to further minimize peristaltic movement. Imaging was carried out on a Leica SP8 multiphoton microscope (Leica Microsystems) with a 20X 1.0 NA water-dipping objective and equipped with a resonant scanner (8,000 kHz) and four external HyD detectors. The excitation wavelength of Chameleon Vision II Ti:Sapphire laser (Coherent) was tuned to 900 nm. After imaging, mice were euthanized by CO_2_ asphyxiation. Images were analyzed with Imaris imaging software (Oxford Instruments).

### Statistics

Statistical significance was calculated using unpaired t test with Welch’s correction for comparing groups of 2 or ANOVA followed by multiple comparisons for comparing groups of 3 or more. Analyses were performed using GraphPad Prism v9.

### Data Availability

All data generated or analyzed during this study are included in this published article (including the supplementary information files).

## Acknowledgements

This work was supported in part by the National Institutes of Health with grants to C.A.H. and B.S. (U01AI163671 and R01AI148249), to B.S. (R01AI112427), to C.A.H. (U01AI160664, R01AI157247), a fellowship to I.S.C. (F30AI169744-01A1), training grant support to B.E.H., J.E.D., and L.A.S. (T32AI007532), J.A.G. (T32AI055400), K.M.O (T32AI055428), a fellowship from EMBO to A.G. (ALTF58-2018), and a fellowship from the Canadian Institutes of Health Research (MFE-176621) and a Postdoctoral Training award from the Fonds de Recherche du Québec – Santé (300355) to R.D.P. B.S. and C.A.H. are supported by the Commonwealth of Pennsylvania.

## Author Contributions

B.E.H., J.A.G., B.S., and C.A.H. conceived of these studies. B.E.H. and J.A.G. conducted experiments with help from I.S.C., K.M.O., R.D.P., M.I.M., and L.A.S. J.E.D., E.N.H., B.A.W. and A.G. engineered transgenic parasites. B.A.W. and A.G. imaged parasite-infected cells. B.A.W. performed multiphoton live imaging. L.A.S., J.H.B., E.J.S., G.Y.B., B.I.M., and D.A.C. bred, maintained, and provided mice for these studies. All authors approved the final manuscript and agree to be accountable for all aspects of the work in ensuring that questions related to the accuracy or integrity of any part of the work are appropriately investigated and resolved.

## References

1. Elliott, D. A. & Clark, D. P. Cryptosporidium parvum Induces Host Cell Actin Accumulation at the Host-Parasite Interface. Infect. Immun. 68, 2315–2322 (2000).

2. Kotloff, K. L. et al. Burden and aetiology of diarrhoeal disease in infants and young children in developing countries (the Global Enteric Multicenter Study, GEMS): a prospective, case-control study. The Lancet 382, 209–222 (2013).

3. Aulagnon, F., Scemla, A., DeWolf, S., Legendre, C. & Zuber, J. Diarrhea After Kidney Transplantation: A New Look at a Frequent Symptom. Transplantation 98, 806–816 (2014).

4. Lanternier, F., et al. Cryptosporidium spp. Infection in Solid Organ Transplantation: The Nationwide “TRANSCRYPTO” Study. Transplantation 101, 826–830 (2017).

5. Wamae, C. N. et al. Cryptosporidiosis in HIV/AIDS Patients in Kenya: Clinical Features, Epidemiology, Molecular Characterization and Antibody Responses. Am. J. Trop. Med. Hyg. 91, 319–328 (2014).

6. McDonald, V. & Bancroft, G. J. Mechanisms of innate and acquired resistance to Cryptosporidium parvum infection in SCID mice. Parasite Immunol. 16, 315–320 (1994).

7. Ungar, B. L., Kao, T. C., Burris, J. A. & Finkelman, F. D. Cryptosporidium infection in an adult mouse model. Independent roles for IFN-gamma and CD4+ T lymphocytes in protective immunity. J. Immunol. 147, 1014–1022 (1991).

8. Sateriale, A., et al. A Genetically Tractable, Natural Mouse Model of Cryptosporidiosis Offers Insights into Host Protective Immunity. Cell Host Microbe 26, 135–146.e5 (2019).

9. Russler-Germain, E. V. et al. Commensal Cryptosporidium colonization elicits a cDC1-dependent Th1 response that promotes intestinal homeostasis and limits other infections. Immunity 54, 2547–2564.e7 (2021).

10. Leav, B. A. et al. An Early Intestinal Mucosal Source of Gamma Interferon Is Associated with Resistance to and Control of Cryptosporidium parvum Infection in Mice. Infect. Immun. 73, 8425–8428 (2005).

11. Ward, H. D. et al. Human CD8+ T Cells Clear Cryptosporidium parvum from Infected Intestinal Epithelial Cells. Am. J. Trop. Med. Hyg. 82, 600–607 (2010).

12. Lantier, L. et al. Intestinal CD103+ Dendritic Cells Are Key Players in the Innate Immune Control of Cryptosporidium parvum Infection in Neonatal Mice. PLoS Pathog. 9, e1003801 (2013).

13. Potiron, L. et al. Batf3-Dependent Intestinal Dendritic Cells Play a Critical Role in the Control of Cryptosporidium parvum Infection. J. Infect. Dis. 219, 925–935 (2019).

14. Bedi, B., McNair, N. N. & Mead, J. R. Dendritic cells play a role in host susceptibility to Cryptosporidium parvum infection. Immunol. Lett. 158, 42–51 (2014).

15. Barakat, F. M., McDonald, V., Di Santo, J. P. & Korbel, D. S. Roles for NK Cells and an NK Cell-Independent Source of Intestinal Gamma Interferon for Innate Immunity to Cryptosporidium parvum Infection. Infect. Immun. 77, 5044–5049 (2009).

16. Choudhry, N., Petry, F., van Rooijen, N. & McDonald, V. A Protective Role for Interleukin 18 in Interferon γ–Mediated Innate Immunity to Cryptosporidium parvum That Is Independent of Natural Killer Cells. J. Infect. Dis. 206, 117–24 (2012).

17. Gullicksrud, J. A. et al. Enterocyte–innate lymphoid cell crosstalk drives early IFN-γ-mediated control of Cryptosporidium. Mucosal Immunol. 15, 362–372 (2022).

18. Wang, H.-C., Zhou, Q., Dragoo, J. & Klein, J. R. Most Murine CD8+ Intestinal Intraepithelial Lymphocytes Are Partially But Not Fully Activated T Cells. J. Immunol. 169, 4717–4722 (2002).

19. Klein, J. R. T-Cell Activation in the Curious World of the Intestinal Intraepithelial Lymphocyte. Immunol. Res. 30, 327–338 (2004).

20. Vandereyken, M., James, O. J. & Swamy, M. Mechanisms of activation of innate-like intraepithelial T lymphocytes. Mucosal Immunol. 13, 721–731 (2020).

21. Tsitsiklis, A., Bangs, D. J. & Robey, E. A. CD8+ T Cell Responses to Toxoplasma gondii: Lessons from a Successful Parasite. Trends Parasitol. 35, 887–898 (2019).

22. Garg, N., Nunes, M. P. & Tarleton, R. L. Delivery by Trypanosoma cruzi of proteins into the MHC class I antigen processing and presentation pathway. J. Immunol. 158, 3293–3302 (1997).

23. Sheridan, B. S. et al. Oral Infection Drives a Distinct Population of Intestinal Resident Memory CD8+ T Cells with Enhanced Protective Function. Immunity 40, 747–757 (2014).

24. Montagna, G. N. et al. Antigen Export during Liver Infection of the Malaria Parasite Augments Protective Immunity. mBio 5, (2014).

25. Dumaine, J. E. et al. The enteric pathogen Cryptosporidium parvum exports proteins into the cytosol of the infected host cell. eLife 10, e70451 (2021).

26. Guérin, A. et al. Cryptosporidium rhoptry effector protein ROP1 injected during invasion targets the host cytoskeletal modulator LMO7. Cell Host Microbe 29, 1407–1420.e5 (2021).

27. Vinayak, S. et al. Genetic modification of the diarrhoeal pathogen Cryptosporidium parvum. Nature 523, 477–480 (2015).

28. Guérin, A. et al. Cryptosporidium uses multiple distinct secretory organelles to interact with and modify its host cell. Cell Host Microbe 31, 650–664.e6 (2023).

29. Gut, J. & Nelson, RG. Cryptosporidium parvum: synchronized excystation in vitro and evaluation of sporozoite infectivity with a new lectin-based assay. J. Eukaryot. Microbiol. 46, 56S–57S (1999).

30. Moran, A. E. et al. T cell receptor signal strength in Treg and iNKT cell development demonstrated by a novel fluorescent reporter mouse. J. Exp. Med. 208, 1279–1289 (2011).

31. Shallberg, L. A. et al. Impact of secondary TCR engagement on the heterogeneity of pathogen-specific CD8+ T cell response during acute and chronic toxoplasmosis. PLOS Pathog. 18, e1010296 (2022).

32. Herndler-Brandstetter, D. et al. KLRG1+ Effector CD8+ T Cells Lose KLRG1, Differentiate into All Memory T Cell Lineages, and Convey Enhanced Protective Immunity. Immunity 48, 716–729.e8 (2018).

33. Harrington, L. E., Janowski, K. M., Oliver, J. R., Zajac, A. J. & Weaver, C. T. Memory CD4 T cells emerge from effector T-cell progenitors. Nature 452, 356–360 (2008).

34. Durai, V. et al. Cryptic activation of an Irf8 enhancer governs cDC1 fate specification. Nat. Immunol. 20, 1161–1173 (2019).

35. Wang, Y. et al. Structural Analyses of a Dominant Cryptosporidium parvum Epitope Presented by H-2K Offer New Options To Combat Cryptosporidiosis. mBio 14, e02666–22 (2023).

36. Burgess, S. L., Gilchrist, C. A., Lynn, T. C. & Petri, W. A. Parasitic Protozoa and Interactions with the Host Intestinal Microbiota. Infect. Immun. 85, e00101–17 (2017).

37. Hand, T. W. et al. Acute Gastrointestinal Infection Induces Long-Lived Microbiota-Specific T Cell Responses. Science 337, 1553–1556 (2012).

38. Lin, Y. H. et al. Small intestine and colon tissue-resident memory CD8+ T cells exhibit molecular heterogeneity and differential dependence on Eomes. Immunity 56, 207–223.e8 (2023).

39. Milner, J. J. et al. Runx3 programs CD8+ T cell residency in non-lymphoid tissues and tumours. Nature 552, 253–257 (2017).

40. Mueller, S. N. & Mackay, L. K. Tissue-resident memory T cells: local specialists in immune defence. Nat. Rev. Immunol. 16, 79–89 (2016).

41. Mackay, L. K. et al. T-box Transcription Factors Combine with the Cytokines TGF-β and IL-15 to Control Tissue-Resident Memory T Cell Fate. Immunity 43, 1101–1111 (2015).

42. Hochheiser, K. et al. Ptpn2 and KLRG1 regulate the generation and function of tissue-resident memory CD8+ T cells in skin. J. Exp. Med. 218, e20200940 (2021).

43. McDole, J. R. et al. Goblet cells deliver luminal antigen to CD103+ dendritic cells in the small intestine. Nature 483, 345–349 (2012).

44. Niess, J. H. CX3CR1-Mediated Dendritic Cell Access to the Intestinal Lumen and Bacterial Clearance. Science 307, 254–258 (2005).

45. Hapfelmeier, S. et al. Microbe sampling by mucosal dendritic cells is a discrete, MyD88-independent stepin ΔinvG S. Typhimurium colitis. J. Exp. Med. 205, 437–450 (2008).

46. Schulz, O. et al. Intestinal CD103+, but not CX3CR1+, antigen sampling cells migrate in lymph and serve classical dendritic cell functions. J. Exp. Med. 206, 3101–3114 (2009).

47. Deets, K. A., Nichols Doyle, R., Rauch, I. & Vance, R. E. Inflammasome activation leads to cDC1-independent cross-priming of CD8 T cells by epithelial cell-derived antigen. eLife 10, e72082 (2021).

48. Sateriale, A., et al. The intestinal parasite Cryptosporidium is controlled by an enterocyte intrinsic inflammasome that depends on NLRP6. Proc. Natl. Acad. Sci. 118, e2007807118 (2021).

49. de Koning-Ward, T. F. et al. A newly discovered protein export machine in malaria parasites. Nature 459, 945–949 (2009).

50. Hakimi, M.-A., Olias, P. & Sibley, L. D. Toxoplasma Effectors Targeting Host Signaling and Transcription. Clin. Microbiol. Rev. 30, 615–645 (2017).

51. Cockburn, I. A. et al. Dendritic Cells and Hepatocytes Use Distinct Pathways to Process Protective Antigen from Plasmodium in vivo. PLoS Pathog. 7, e1001318 (2011).

52. Gregg, B. et al. Subcellular Antigen Location Influences T-Cell Activation during Acute Infection with Toxoplasma gondii. PLoS ONE 6, e22936 (2011).

53. Marino, N. D. et al. Identification of a novel protein complex essential for effector translocation across the parasitophorous vacuole membrane of Toxoplasma gondii. PLOS Pathog. 14, e1006828 (2018).

54. Curt-Varesano, A., Braun, L., Ranquet, C., Hakimi, M.-A. & Bougdour, A. The aspartyl protease TgASP5 mediates the export of the Toxoplasma GRA16 and GRA24 effectors into host cells: TgASP5 is essential for GRA16 and GRA24 export. Cell. Microbiol. 18, 151–167 (2016).

55. Pepper, M., Dzierszinski, F., Crawford, A., Hunter, C. A. & Roos, D. Development of a System To Study CD4-T-Cell Responses to Transgenic Ovalbumin-Expressing Toxoplasma gondii during Toxoplasmosis. INFECT IMMUN 72, 7 (2004).

56. Lin, J.-S., Szaba, F. M., Kummer, L. W., Chromy, B. A. & Smiley, S. T. Yersinia pestis YopE Contains a Dominant CD8 T Cell Epitope that Confers Protection in a Mouse Model of Pneumonic Plague. J. Immunol. 187, 897–904 (2011).

57. Sateriale, A., Pawlowic, M., Vinayak, S., Brooks, C. & Striepen, B. Genetic Manipulation of Cryptosporidium parvum with CRISPR/Cas9. Methods Mol. Biol. Clifton NJ 2052, 219–228 (2020).

58. Manjunatha, U. H. et al. A Cryptosporidium PI(4)K inhibitor is a drug candidate for cryptosporidiosis. Nature 546, 376–380 (2017).

