## Supplemental Information for "Dendritic cell-mediated responses to secreted *Cryptosporidium* effectors are required for parasite-specific CD8^+^ T cell responses"

Supplemental Figure 1.

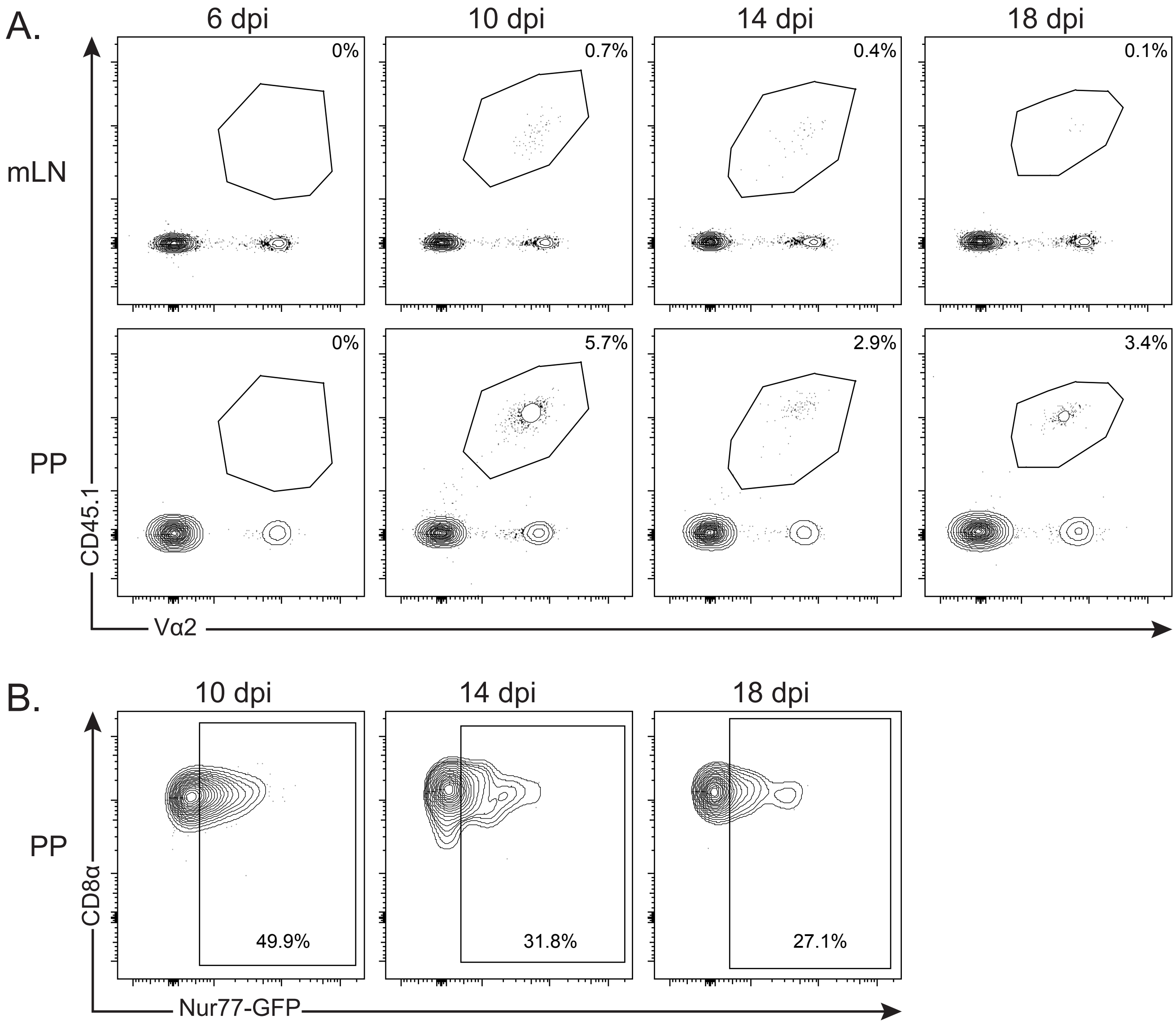


Supplemental Figure 1. (Refers to Figure 2). **A**. Representative flow plots display OT-I cells in mLN and PP from experiments performed in Fig. 2A/B. **B**. Representative flow plots of Nur77-GFP OT-I cells in PP, from experiments performed in Fig. 2E.

### Supplemental Figure 2.

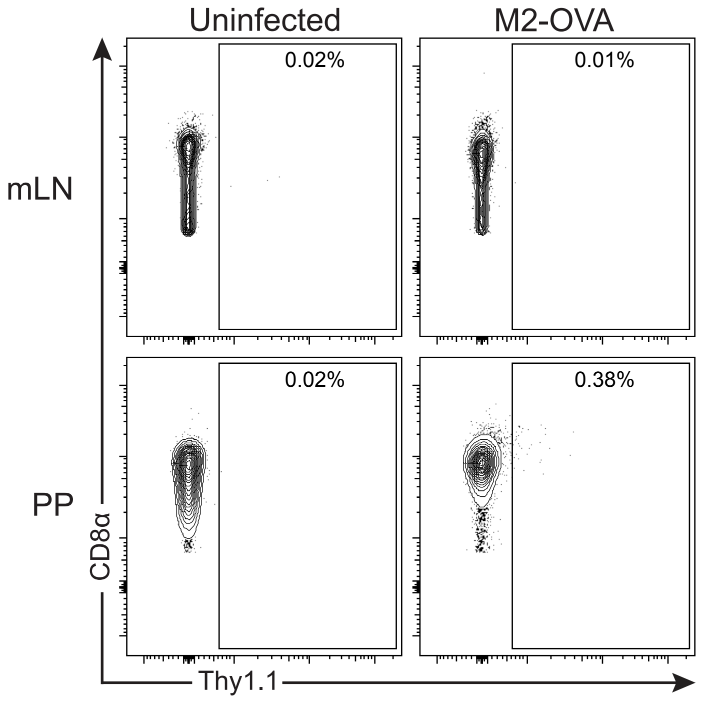


Supplemental Figure 2. (Refers to Figure 4). Representative flow plots show CD8⁺ T cell Thy1.1 expression in mLN and PP from experiments performed in Fig. 4A.

Supplemental Figure 3.

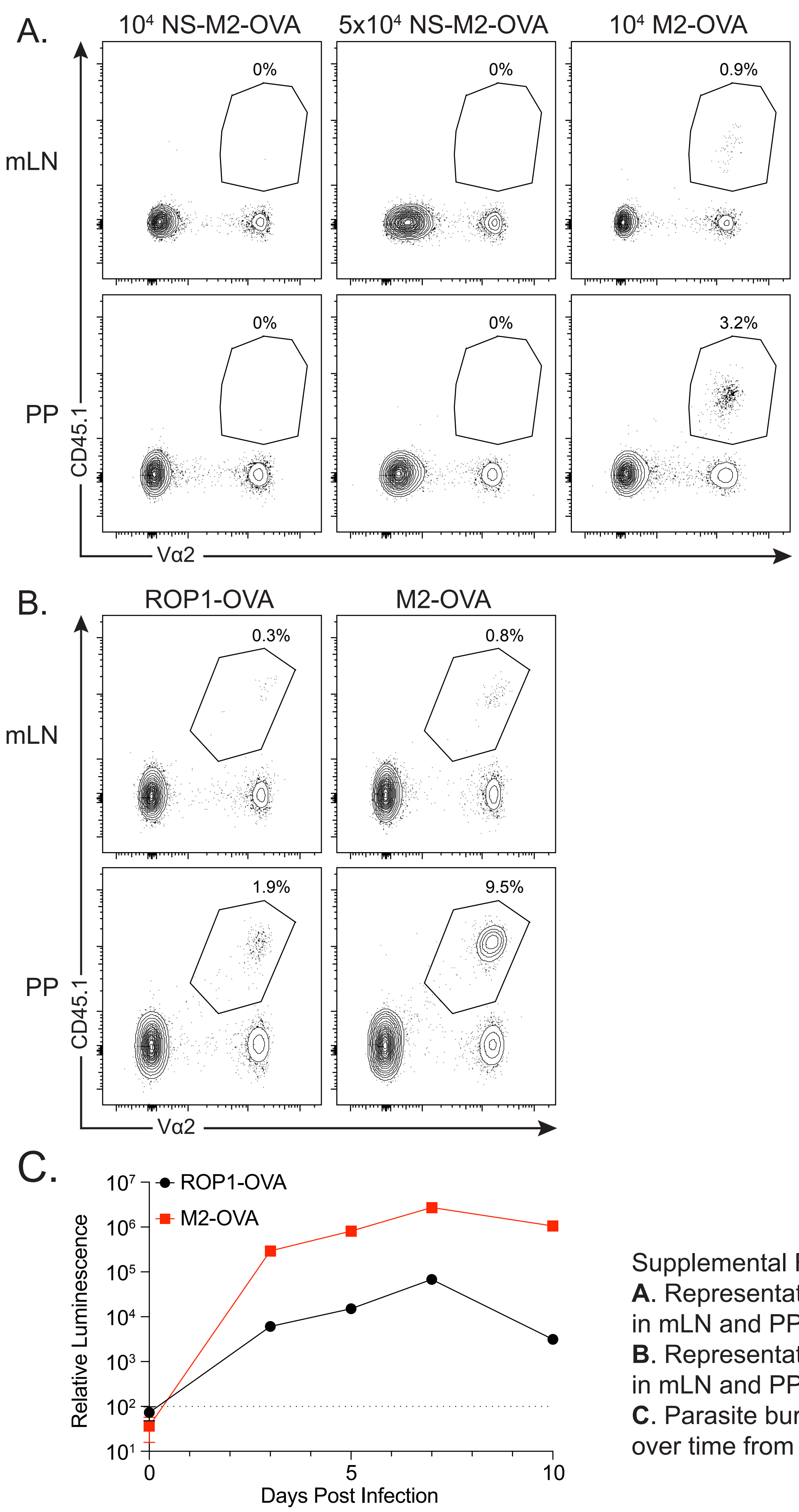


Supplemental Figure 3. (Refers to Figure 5). **A**. Representative flow plots show OT-I cells in mLN and PP from experiments in Fig. 5C. **B**. Representative flow plots show OT-I cells in mLN and PP from experiments in Fig. 5D. **C**. Parasite burdens (relative luminescence) over time from mice infected in Fig. 5D.

### Supplemental Figure 4.

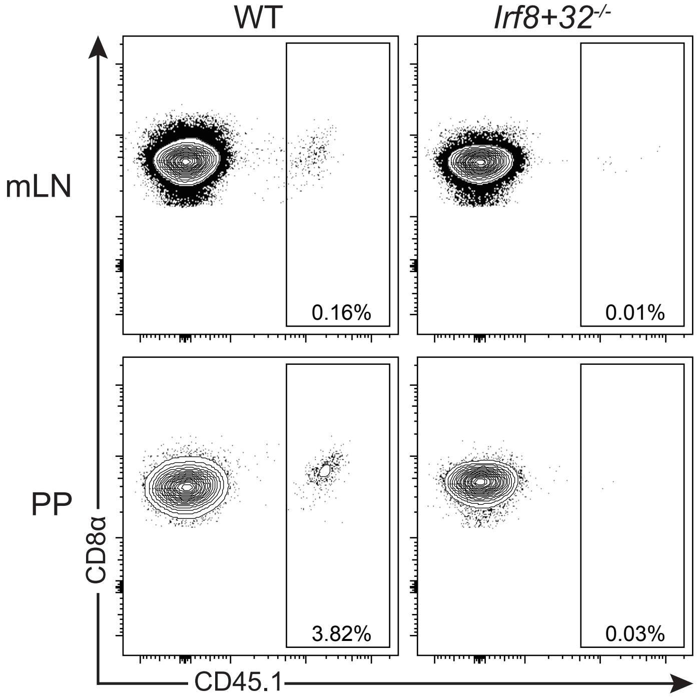


Supplemental Figure 4. (Refers to Figure 6). Representative flow plots show OT-I cells in mLN and PP from experiments in Fig. 6B.

| Supplemental Table 1. List of Primers for plasmid construction | |
|---|---|
| Use | Sequence |
| Linearizing p3XHA_nLuc-Neo for SIINFEKL insertion | taacccgggatgcatcttca |
| | ggcataatctggaacatcgtaagg |
| SIINFEKL sequence for Gibson assembly | ccttacgatgttccagattatgccagtataatcaactttgaaaaactg |
| | tgaagatgcatcccgggttacagtttttcaaagttgattatact |
| Repair template amplification M2-OVA | aaaaagaaaaagagaagaggggaaaagatgatcttagaggactattaagaggattatccggttcaagtaaagacatactacacgacttagagaaaattggaagtggaggacgggaattc |
| | taattatcttgttaataacacatttgaatagtttatacttcacataaccaaattaagataaaaagaaaaacttaatcgatactatcctacacg |
| M2 targeting guide | gttgaggattatccggttcaagta |
| | aaactacttgaaccggataatcct |
| Repair template amplification ROP1-OVA | tataaatcacctgaattccgaaaatggaagtggaggacgggaattc |
| | gaatttgggtattttctcgtctcaccgcgtttaaactgattggtacta |
| ROP1 targetting guide | gttggactaaaaataattgtttcaa |
| | aaacttgaaacaattatttttagtc |
| Linearizing p3XHA_nLuc-Neo_M2 for tdTom insertion | aattggttcgtggcgtgtagga |
| | gaagaattcgtcaagaagacgatagaag |
| tdTom amplification for Gibson assembly | gccttctatcgtcttcttgacgaattcttcggtgagggcaggggtagattgttgacttgcggtgacgttgaggagaaccccggcccgatggtgagtaagggcgag |
| | tcacttatacagctcatccatgc |
| Repair template NS-M2-OVA | tagctttttgccacagcgacaaatagttttgatttcagtaagtttatcaccatagctgcgccaaattttgc |
| | tccagtactatgctatggtttgagaacagactttaagggaaatttatttgatggggaaactaaatatactgaaattcggt |
| TK targetting guide for NS-M2-OVA insertion | gttggaaggaagtaaatacttattagca |
| | aaactgctaataagtatttacttc |
